# The inner nuclear membrane protein SUN1 regulates cullin-3 neddylation to maintain insulin signaling

**DOI:** 10.64898/2026.04.16.718478

**Authors:** Kapil K. Upadhyay, Yunshu Yang, Aren Shah, Venkatesha Basrur, Alexey I. Nesvizhskii, Graham F. Brady

**Author notes:** **Corresponding Author**: University of Michigan Medical School, Division of Gastroenterology and Hepatology, Department of Internal Medicine, 3912 Taubman Center, 1500 East Medical Center Drive, Ann Arbor, MI, USA 48109-5362. **Conflict of interest statement:** The authors declare no conflicts of interest. **Author Contributions:** K.K.U. and G.F.B. conceived and designed the study; K.K.U., A.S., Y.Y., and V.B. performed the *in vitro* experiments and mass spectrometric analysis; K.K.U., A.S., V.B., A.N., and G.F.B. analyzed the data; K.K.U. and G.F.B. wrote the manuscript; all authors critically reviewed and edited the manuscript.

## Abstract

Metabolic dysfunction-associated steatotic liver disease (MASLD) is the most prevalent chronic liver disease and strongly linked to obesity and insulin resistance. We previously reported that the common nuclear envelope variant rs6461378 (g.842031C>T; *SUN1* H118Y) associated with MASLD and related traits including insulin resistance. To gain insight into how wild-type (WT) and H118Y SUN1 might differentially impact insulin signaling, we performed affinity purification-mass spectrometry (AP-MS) in human liver-derived cells stably expressing WT or H118Y SUN1. Unbiased AP-MS revealed a novel SUN1-CUL3 interaction, with comparative analysis showing that WT SUN1 interacted robustly with CUL3, while CUL3 interaction was markedly diminished with H118Y SUN1. Cells in which *SUN1* was silenced via siRNA, or in which H118Y SUN1 was ectopically expressed, showed increased CUL3 neddylation, which is required for cullin RING ligase (CRL)-mediated ubiquitination of insulin receptor substrate (IRS) proteins. Inhibition of neddylation restored IRS-1 levels and insulin signaling in H118Y SUN1-expressing cells. Together, our findings provide a potential mechanism of H118Y SUN1-driven insulin resistance and a viable therapeutic approach for its reversal.

## Introduction

Metabolic dysfunction–associated steatotic liver disease (MASLD) is now estimated to affect nearly one-third of the population worldwide, with a rising incidence that parallels increases in obesity, metabolic syndrome, insulin resistance, and type 2 diabetes (Cotter & Rinella, 2020; Rinella *et al*, 2023; Samuel & Shulman, 2018). In patients with MASLD, the presence or absence of insulin resistance is a key determinant of both MASLD progression, with accompanying long-term risks of cirrhosis and hepatocellular carcinoma, as well as overall cardiometabolic risk, though the molecular mechanisms of insulin resistance are incompletely understood (Bo *et al*, 2024; Loomba *et al*, 2021; Steinberg *et al*, 2025).

Rare variants in genes encoding nuclear envelope and lamina proteins, such as *LMNA* (encoding A-type lamins), are well-established causes of monogenic metabolic diseases, including familial partial lipodystrophy, characterized by severe insulin resistance, aberrant adipose distribution, and early-onset MASLD. Recent evidence from a combination of human genetics and mouse models suggests, though, that nuclear envelope proteins play key roles in protecting from, or predisposing to, MASLD and metabolic disease more broadly (Brady *et al*, 2018; Chen *et al*, 2023; Emdin *et al*, 2021; Kwan *et al*, 2017; Shin *et al*, 2019). Extending this paradigm, we recently identified a common coding variant, rs6461378 (g.842031C>T; H118Y) in *SUN1*, encoding the inner nuclear membrane protein Sad1 and unc84 domain containing 1 (SUN1), in association with hepatic steatosis, histologically defined MASLD, and related metabolic traits, including insulin resistance (Upadhyay *et al*, 2023). *In vitro* proof-of-concept studies confirmed rs6461378-T as a functional variant with a potential contributory role in insulin resistance and MASLD; expression of H118Y SUN1 in Huh7 or HepG2 cells caused lipid droplet accumulation and reduced AKT phosphorylation in response to insulin compared to wild-type (WT) SUN1. However, the precise molecular mechanisms linking *SUN1* H118Y to insulin resistance are not yet known.

With its partner SUN2 and nesprins, SUN1 forms part of the <u>Li</u>nker of <u>N</u>ucleoskeleton and <u>C</u>ytoskeleton (LINC) complex, which physically links the cytoskeleton to the genome via its interaction with the nuclear lamina in all nucleated metazoan cells. Despite physical linkage to the genome via the lamina, however, recent data from animal models, including mouse and zebrafish, indicate that SUN1 – unlike nuclear lamina proteins – likely does not directly regulate chromatin accessibility or gene transcription. Given the location of the residue impacted by rs6461378-T (His118, located within the SUN1 intranuclear amino-terminal domain that is required for interaction with lamins and other intranuclear proteins), we hypothesized that differential intranuclear protein-protein interactions might mediate the distinct effects of WT and H118Y SUN1 on insulin signaling. Here we report that, via unbiased affinity purification-mass spectrometry (AP-MS), we identified a novel SUN1-cullin-3 (CUL3) interaction that regulates insulin signaling in human cells. Compared to WT SUN1, co-precipitation of endogenous CUL3 was dramatically reduced by the presence of the H118Y variant. Furthermore, WT SUN1 reduced post-translational neddylation and activation of CUL3, with increased activation of CUL3-containing cullin E3 RING ubiquitin ligase (CRL3) complexes in H118Y SUN1-expressing cells, accompanied by reduced levels of CRL3 degradation targets including insulin receptor substrate-1 (IRS-1). Lastly, insulin signaling could be restored in H118Y SUN1-expressing cells by pharmacologic inhibition of CUL3 neddylation. These findings define a novel SUN1–CUL3 regulatory mechanism in insulin signaling and position the SUN1-CUL3 interaction as a potential therapeutic target in MASLD and insulin resistance.

## Materials and Methods

### Cell Culture and Treatments

Huh7 cells from the Japanese Collection of Research Bioresources (JCRB) were cultured in Dulbecco’s Modified Eagle Medium (DMEM; Gibco) containing glucose but lacking L-glutamine, supplemented with 10% heat-inactivated fetal bovine serum (FBS; Thermo Fisher Scientific, A3840202, USA). Cells were maintained at 37°C in a humidified incubator with 5% CO₂. For transfection experiments, cells were seeded to achieve approximately 60–70% confluency one day prior to transfection. To inhibit neddylation, cells were treated with either 1 μM MLN4924 (Sigma-Aldrich, St. Louis, MO), a pan-neddylation inhibitor, or with a CUL3-specific neddylation inhibitor (DI-1859, Med Chem Express HY-145733) at the indicated concentrations for 16 hours to achieve selective inhibition of CUL3 neddylation. Where indicated, recombinant human insulin (Sigma-Aldrich, catalog #91077C) was added to serum-free culture medium prior to lysis, as described (Upadhyay *et al*., 2023).

### Recombinant lentivirus and lentiviral transduction of Huh7 cells

To generate Huh7 cell lines stably expressing either WT or H118Y SUN1 with a carboxy-terminal mCherry tag, the full-length coding sequences for WT SUN1 or H118Y SUN1 were PCR-amplified from previously described plasmids (Upadhyay *et al*., 2023) using the oligonucleotide primers (Forward primer: 5′-GCTTGATTGATTCGAGCTAGCATCTGCCGCCGCGATCGC-3′; Reverse primer: 5′-AACCCGGGAATGGGCCCAACTCGAGAAGGACAGGGAAGGGAGCAGTG-3′) containing an XhoI and BmtI site for each end. The resulting PCR products were subcloned into the pLentiLox-CAG-MCS-EFS-Puro A vector (obtained from the University of Michigan Vector Core) containing a puromycin resistance cassette using the NEB HiFi cloning kit (New England Biolabs, Ipswich, MA). Lentiviral particles were produced by the University of Michigan Vector Core in human embryonic kidney (HEK) 293T cells using the VSV-G packaging system. Huh7 cells were then transduced with 1X lentiviral supernatant (a titer was not determined) in the presence of polybrene (10 μg/mL) and incubated at 37°C, 5% CO₂ for 48 hours. Following transduction, stably expressing cells were selected in culture medium containing puromycin (8 μg/mL). Selection was continued for two weeks, by which time all control, untransduced Huh7 cells maintained in parallel and also treated with 8 μg/mL puromycin had died. Selected, puromycin-resistant, transduced Huh7 cells were maintained in puromycin-containing medium (4 μg/mL) for subsequent experiments.

### Plasmids, siRNA, and Transfection

For transient expression studies, a plasmid encoding human WT or H118Y SUN1-mCherry (Upadhyay *et al*., 2023) was transfected into Huh7 cells with Lipofectamine-2000 (Life Technologies, Carlsbad, CA) according to the manufacturer’s protocol. For transient silencing of endogenous *SUN1* in Huh7 cells, cells were transfected with a *SUN1*-targeting siRNA (Sigma-Aldrich; siRNA ID: SASI_Hs01_00032811) using Lipofectamine RNAiMAX (Thermo Fisher) according to the manufacturer’s protocol. A universal negative control siRNA (Sigma Aldrich SIC001) was used as the control for all RNA interference experiments.

### mCherry-Affinity Immunoprecipitation

Huh7 cells stably expressing WT or H118Y SUN1-mCherry were cultured in DMEM supplemented with 10% FBS. Cells were seeded in 15-cm culture dishes and allowed to reach confluency (∼48 hours). Upon reaching 100% confluency, cells were washed twice with sterile phosphate-buffered saline (PBS) and harvested by scraping into ice-cold radioimmunoprecipitation assay (RIPA) buffer (Thermo Fisher Scientific) supplemented with protease inhibitors (Sigma-Aldrich) and phosphatase inhibitors (Thermo Fisher Scientific). Cell lysates were clarified by centrifugation at 14,000 × g for 10 minutes at 4°C, and the supernatants were collected for affinity purification. For immunoprecipitation, DynaGreen™ magnetic beads (Thermo Fisher Scientific) were resuspended by vortexing, and 50 μL aliquots were transferred into clean microcentrifuge tubes. Beads were magnetically separated, washed, and incubated with 20 µg of anti-mCherry antibody (Thermo Fisher Scientific) in 500 μL PBS for 1 hour at room temperature with continuous rotation. After antibody coupling, the beads were washed with PBS and resuspended in 500 μL PBS for IP. For each IP, 4 mg of clarified total protein lysate (in 1,000 μL volume) was incubated with antibody-coupled beads for 1 hour at room temperature with rotation. After incubation, beads were washed three times with 500 μL PBS using magnetic separation. The final bead complexes were collected and stored at −80°C until being subjected to mass spectrometry analysis.

### SUN1-interacting protein identification via LC-Tandem MS

Magnetic beads bound to SUN1-mCherry and associated proteins were thawed and resuspended in 50 µL of 0.1 M ammonium bicarbonate buffer (pH∼8). Cysteines were reduced by adding 50 µL of 10 mM DTT and incubating at 45 °C for 30 min. Samples were cooled to room temperature, and alkylation of cysteines was achieved by incubating with 65 mM 2-Chloroacetamide, under darkness, for 30 min at room temperature. An overnight digestion with 1 µg sequencing grade modified trypsin was carried out at 37 °C with constant shaking in a Thermomixer. The digestion was quenched by acidification, and the resulting peptides were desalted via Sep-Pak C18 cartridges (Waters) according to the manufacturer’s protocol. Peptides were vacuum-dried and resuspended in 9 µL of 0.1% formic acid/2% acetonitrile. A 2 µL aliquot was loaded onto a nano-capillary reverse-phase column (Acclaim PepMap C18, 2 µm, 50 cm, Thermo Scientific) and resolved at a flow rate of 300 nL/min using a 180-minute gradient of 0.1% formic acid in water (Buffer A) and 0.1% formic acid in 95% acetonitrile (Buffer B). The gradient profile was: 2–22% B over 110 min, 22–40% B over 25 min, and 40–90% B over 5 min, followed by a 5-minute wash at 90% B and a 30-minute re-equilibration. Eluted peptides were analyzed on a Q Exactive HF mass spectrometer (Thermo Scientific) equipped with an Easy-Spray source. MS1 scans were acquired at 60K resolution (AGC target=3×10^6^; max IT=50 ms). Data-dependent MS/MS spectra were acquired using a 3-second ‘Top Speed’ method via HCD fragmentation (NCE ∼28%; 15K resolution; AGC target 1×10^5^; max IT 45 ms). Proteins were then identified by searching the MS/MS data against the human protein database using Proteome Discoverer (v3.0, Thermo Scientific). Search parameters included MS1 mass tolerance of 10 ppm and fragment tolerance of 0.1 Da; two missed cleavages were allowed; carbamidomethylation of cysteine was considered fixed modification, and oxidation of methionine, deamidation of asparagine and glutamine were considered as potential modifications. False discovery rate (FDR) was determined using Percolator, and proteins/peptides with FDR ≤1% were retained for further analysis.

### Western Blotting

Cells were lysed in ice-cold RIPA buffer (Thermo Fisher) supplemented with 1X protease and phosphatase inhibitor cocktails (Thermo Fisher). Lysates were clarified by centrifugation at 14,000 × g for 10 minutes at 4 °C. Protein concentrations were determined by bicinchoninic acid (BCA) assay (Thermo Fisher). Equal amounts of protein were denatured in the presence of SDS and 2% β-mercaptoethanol, then separated on 4–12% Novex Tris-Glycine gels (Invitrogen) via SDS-PAGE. Proteins were transferred onto polyvinylidene fluoride (PVDF) membranes (Bio-Rad, Hercules, CA) for immunoblotting. Detection was achieved through enhanced chemiluminescence (ECL), and images were captured using a ChemiDoc imaging system. Where indicated, band intensities were quantified by densitometry using ImageJ software (version 1.53t).

### Antibodies

Antibodies used in this study were: anti-mCherry (Thermo Fisher PA5-34974, 1:1000); anti-AKT phospho-Ser473 (Cell Signaling 4060, 1:1000); anti-AKT total (Cell Signaling 4691, 1:1000); anti-IRS1 (Cell Signaling 3407, 1:1000); anti-IRS1 phospho-Tyr612 (Thermo Fisher 44-816G, 1:1000); β-actin (Cell Signaling 4970, 1:1000); anti-CUL3 (Cell Signaling 10450, 1:1000); anti-SUN1 (Merck Millipore ABT273, 1:800).

### Statistical analysis

Unless otherwise specified, for continuous data the two-tailed Student’s *t*-test or the Mann-Whitney U test was used to assess statistical significance. Statistical analyses were performed using GraphPad Prism version 10.

## Results

### Unbiased proteomics identifies CUL3 as a novel SUN1-interacting protein, with impaired interaction with H118Y SUN1

We previously reported that H118Y SUN1 reduced insulin signaling to AKT in human liver-derived cell lines (Upadhyay *et al*., 2023), though the mechanisms underlying this dampened insulin signaling remained unclear. Given that His118 lies within SUN1’s nucleoplasmic region and that recent data suggest SUN1 may not directly regulate chromatin accessibility or gene transcription (Buglak *et al*, 2023), we hypothesized that the H118Y variant of SUN1 might disrupt interactions with specific intranuclear protein partners. To identify interaction partners through which SUN1 might regulate insulin signaling pathways in ways that differ between WT and H118Y SUN1, we generated Huh7 cells stably expressing either WT or H118Y SUN1 with a carboxy-terminal mCherry tag using a lentiviral approach. These cells stably express exogenous mCherry-tagged SUN1 at levels similar to endogenous SUN1 (**Fig.1**). To discover intranuclear SUN1-interacting proteins that differ between WT and H118Y SUN1, stably expressed WT and H118Y SUN1-mCherry were separately immunoprecipitated using an anti-mCherry antibody, and the immunoprecipitates were subjected to proteomic analysis via mass spectrometry in triplicate (**Fig.2A**). Visualization of the top differentially interacting proteins via volcano plot revealed a small cluster of proteins that were most preferentially found in the WT SUN1 immunoprecipitates compared to H118Y SUN1 (**Fig.2B**). Examining this group (**Table S1**; all *P*<10^-15^), we prioritized CUL3 for further validation for two reasons: (1) it had the greatest number of unique peptides among this top group (providing high confidence in its identification) and (2) this protein was reported to regulate hepatic insulin signaling and liver injury (Chen *et al*, 2022; Gu *et al*, 2024). Preferential association of CUL3 with WT SUN1 was confirmed via co-precipitation and immunoblot; stably expressed WT SUN1, but not H118Y SUN1, robustly precipitated endogenous CUL3 from Huh7 cell lysates (**Fig.3**).

**Fig. 1.**
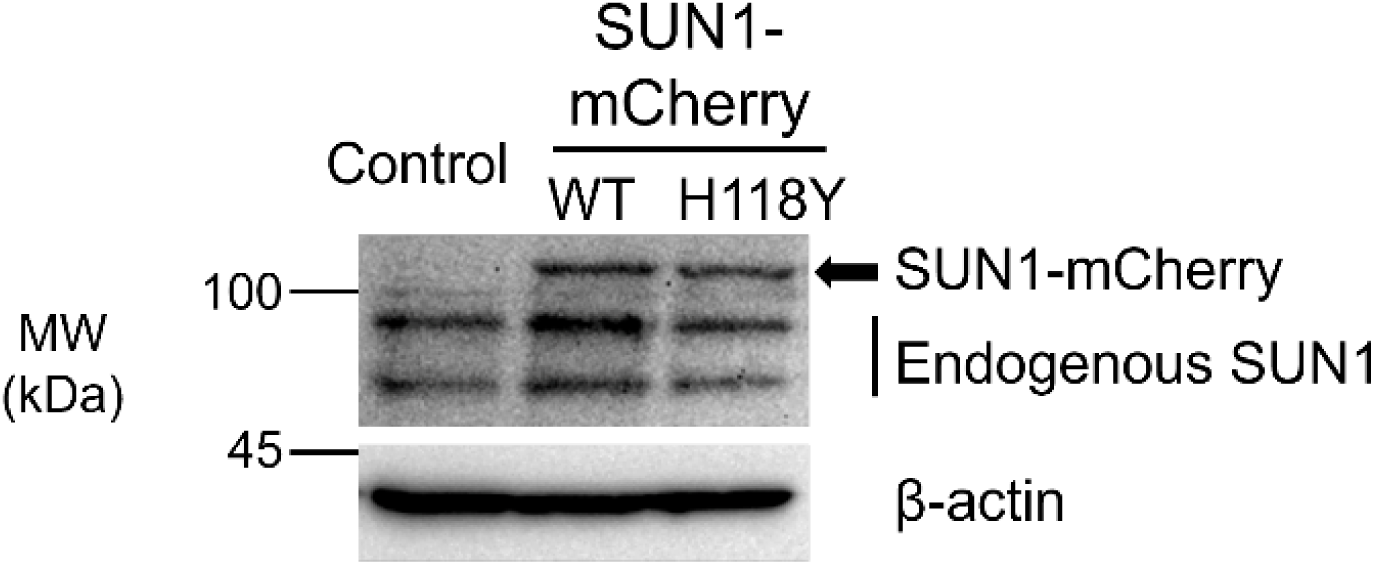
Stable SUN1-expressing Huh7 cells, generated via lentiviral transduction, express SUN1-mCherry at physiologic levels. Exogenous SUN1-mCherry and endogenous SUN1 were detected via anti-SUN1 immunoblot. MW, molecular weight. Results shown are representative of n>3 independent experiments.

**Fig. 2.**
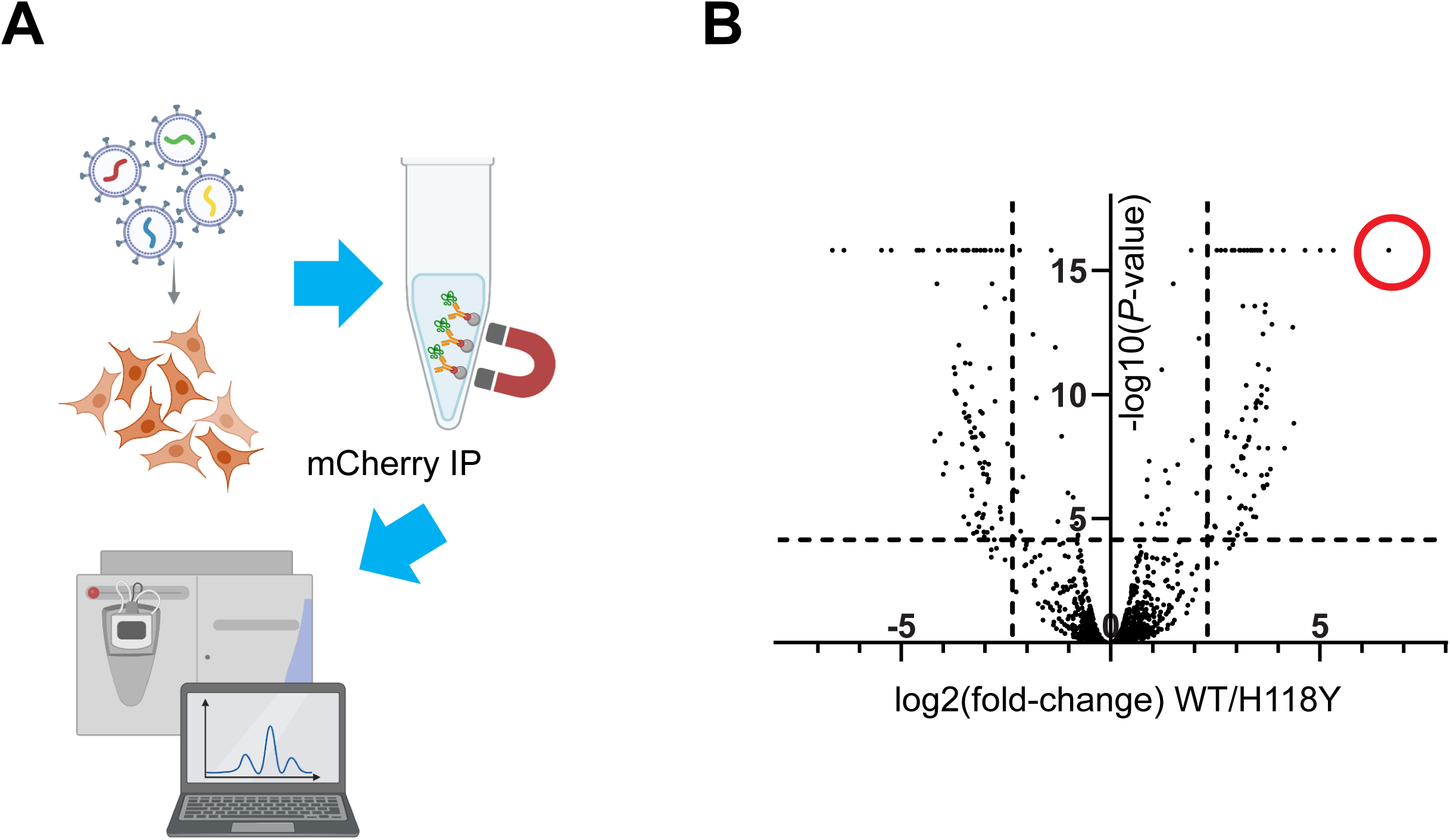
SUN1 affinity purification mass spectrometry (AP-MS) schematic and results. *A*, schematic of lentiviral transduction of Huh7 cells, co-precipitation of SUN1-mCherry with associated proteins, and mass spectrometry to identify differential WT and H118Y SUN1-interacting proteins. *B*, volcano plot of differential WT versus H118Y SUN1-interacting proteins based on n=3 replicate AP-MS experiments; log_2_ fold change >0 indicates WT > H118Y SUN1, while log_2_ fold change <0 indicates H118Y > WT SUN1. The circled cluster at top right includes CUL3 and represents the maximally WT SUN1-enriched set of proteins (with maximum fold change set to 100; *P* ∼ 10^-15^).

**Fig. 3.**
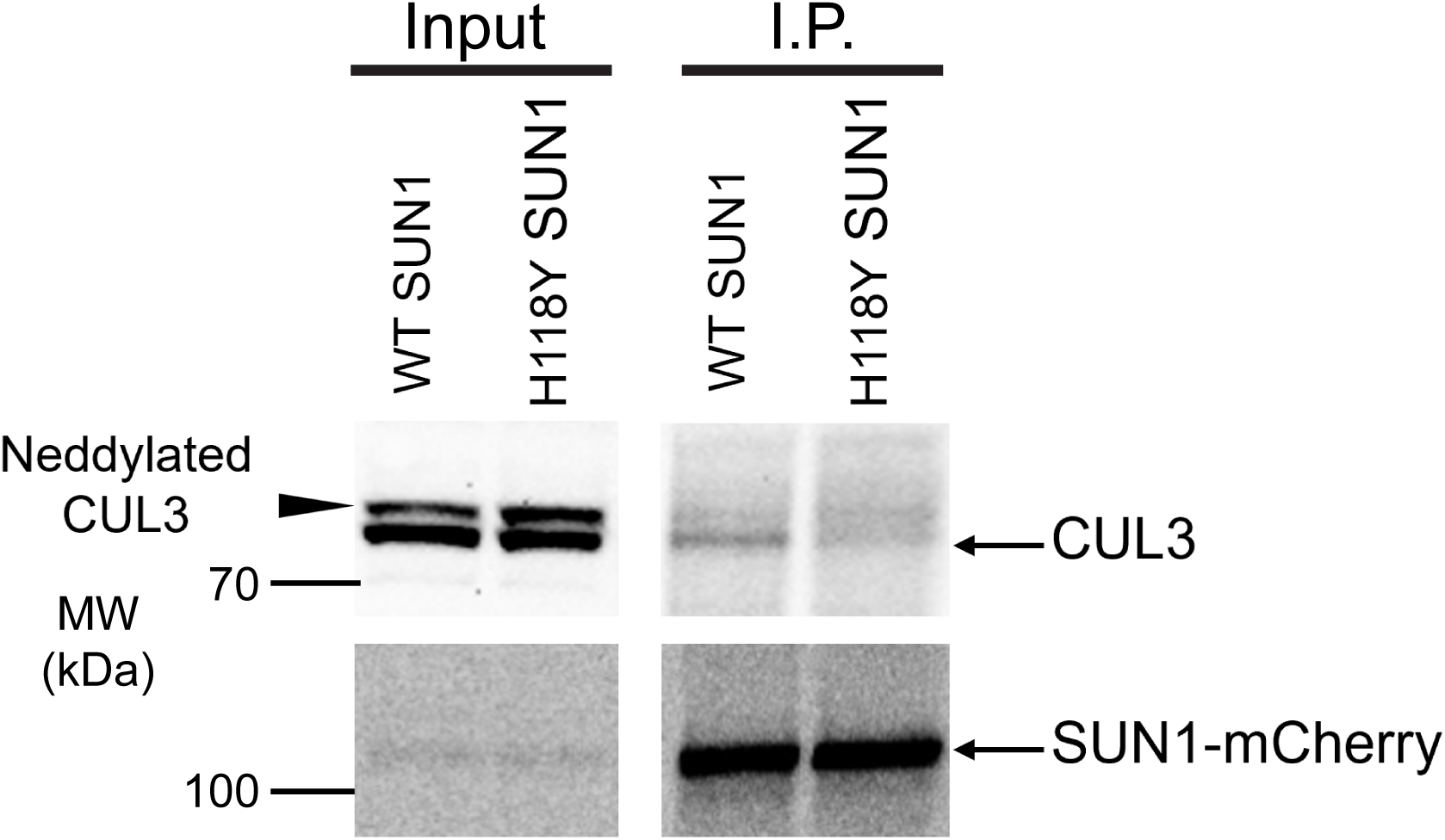
Wild-type (WT) SUN1 co-precipitates endogenous CUL3 to a greater degree than H118Y SUN1 in Huh7 cells. Cells stably expressing SUN1-mCherry were lysed in RIPA buffer followed by mCherry IP, SDS-PAGE, and immunoblot. Native CUL3 is indicated by an arrow, while neddylated CUL3 (higher band) is indicated via arrowhead. Results shown are representative of n=3 independent experiments. MW, molecular weight.

### SUN1 regulates CUL3 neddylation and activation of CUL3-containing RING E3 ubiquitin ligase

In the course of the confirmatory SUN1-CUL3 co-precipitation experiments, it became apparent that a higher molecular weight, slower-migrating CUL3 band was modestly but consistently stronger in the H118Y SUN1 cell lysates compared to WT SUN1 (see input lanes, Fig.3). This higher band recognized by the anti-CUL3 antibody has been reported to represent neddylated CUL3 (Wu *et al*, 2005); activation of CUL3-containing RING E3 ubiquitin ligase (CRL) complexes requires covalent attachment of the ubiquitin-like protein neural precursor cell expressed, developmentally down-regulated 8 (NEDD8) to CUL3 at Lys 712 by a neddylating enzyme of the DCN1-like family (Kurz *et al*, 2008; Lo & Hannink, 2006), with the neddylation status of CUL3 dynamically regulated and balanced via deneddylation by the COP9 signalosome (Fischer *et al*, 2011; Lyapina *et al*, 2001). However, to confirm that this band represented neddylated CUL3 in our system, we treated cells with a neddylation inhibitor (MLN4924) and observed the disappearance of this band, confirming that this band indeed represented neddylated CUL3 (**Fig.S1**).

Given that neddylation of CUL3 is required for activation of CUL3-containing CRL complexes (CRL3), that CRL3 is known to regulate insulin signaling by targeting insulin receptor substrate-1 (IRS-1) for ubiquitination and degradation in hepatocytes (Chen *et al*., 2022), and that CUL3 was previously reported to be sequestered in its inactive, non-neddylated form at the inner nuclear membrane (Mathew *et al*, 2012), we hypothesized that control of CUL3 neddylation might be a mechanism by which SUN1 regulates insulin signaling. To first exclude the possibility that the observed difference in CUL3 neddylation between WT and H118Y SUN1-expressing cells could be due to a SUN1-independent phenomenon related to lentiviral transduction, we transfected parental Huh7 cells with transient expression constructs encoding WT and H118Y SUN1-mCherry: we found that neddylated CUL3 was nearly undetectable in Huh7 cells transfected with WT SUN1-mCherry, while ectopic expression of H118Y SUN1 had no discernible effect (**Fig.4A**). Supporting the idea that restriction of CUL3 neddylation is a physiologic function of SUN1, we found that silencing of endogenous *SUN1* via siRNA in Huh7 cells increased CUL3 neddylation (**Fig.4B**) and reduced IRS-1 levels (**Fig.4C**). Taken together, these data support a model in which WT SUN1 – but not H118Y SUN1, which is apparently not able to efficiently bind to CUL3 – restricts CUL3 neddylation and CRL3 activity to maintain cellular IRS-1 levels.

**Fig. 4.**
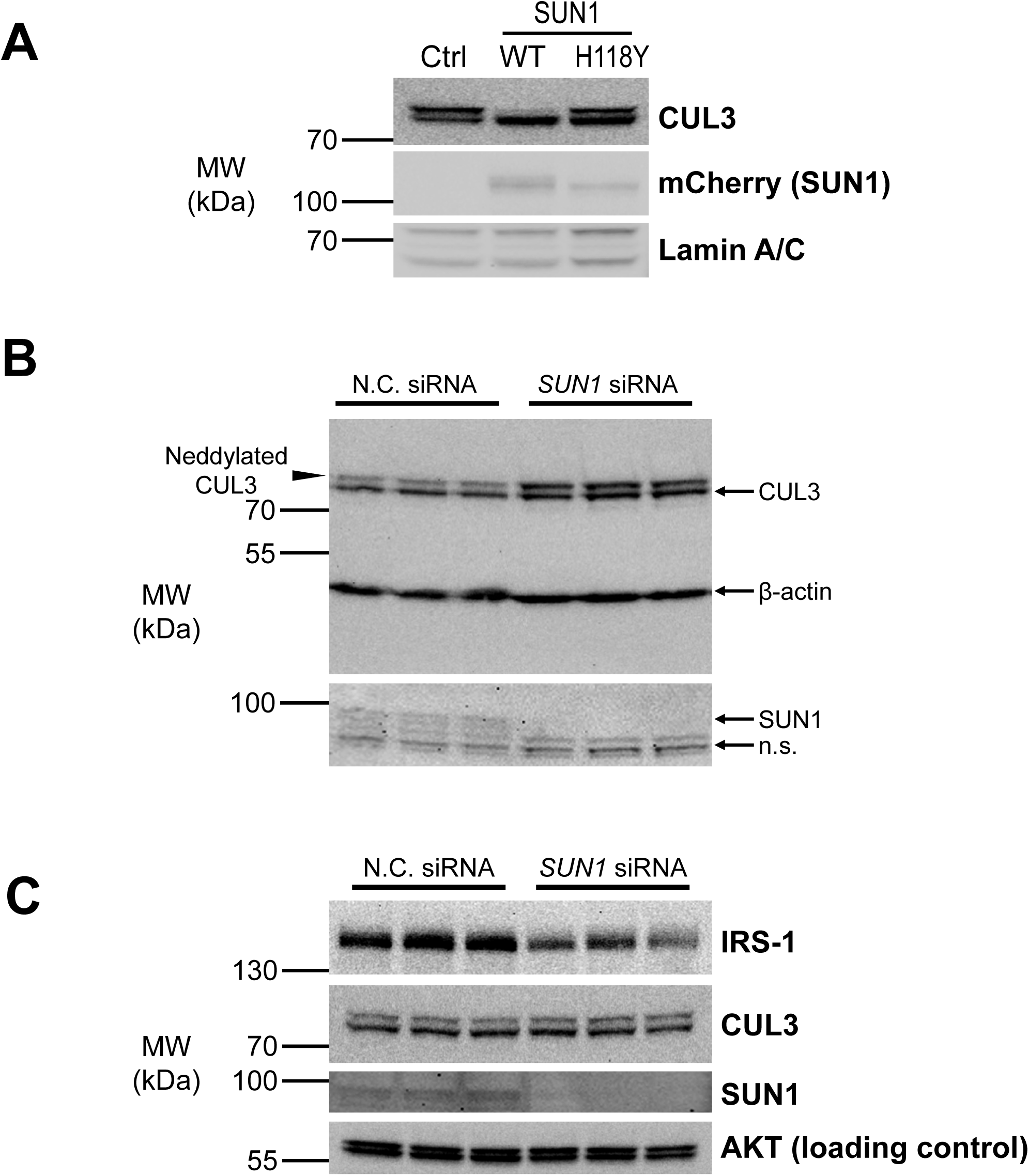
SUN1 regulates CUL3 neddylation and maintains cellular IRS-1 levels. *A*, SUN1-mCherry was transiently expressed in Huh7 cells for 48h prior to lysis and detection of neddylated and total CUL3 with anti-CUL3 antibody. Control, untransfected Huh7 cells are shown in lane 1 for comparison and labeled “Ctrl.” *B* and *C*: Huh7 cells were treated with si*SUN1* oligos or a negative control (N.C.) oligo in triplicate, followed by lysis, SDS-PAGE, and immunoblot for assessment of CUL3 neddylation (part *B*) or IRS-1 levels (part *C*). In *B*, native CUL3 is indicated by an arrow, while neddylated CUL3 (higher band) is indicated via arrowhead. n.s., non-specific bands. Results shown are representative of n=3 independent experiments.

### Inhibition of CUL3 neddylation restores insulin signaling in H118Y SUN1-expressing cells

Given that we previously showed that H118Y SUN1 reduced insulin signaling to AKT in liver-derived cells, we next asked whether this blunted insulin signaling could be due to reduced cellular IRS-1 levels and reversible via pharmacologic inhibition of CUL3 neddylation. To determine whether inhibition of cullin neddylation could rescue insulin signaling in H118Y SUN1-expressing cells, we pre-treated cells with the pan-neddylation inhibitor MLN4924 prior to insulin treatment. Compared to vehicle, pre-treatment with MLN4924 restored IRS-1 levels and increased insulin-induced AKT Ser473 phosphorylation in H118Y SUN1-expressing cells, with both IRS-1 expression and AKT phosphorylation reaching the levels seen in WT SUN1-expressing cells (**Fig.5A**). Thus, the effects of H118Y SUN1 on cellular IRS-1 levels and insulin signaling to AKT were completely reversed by inhibiting CUL3 neddylation and CRL3 activation. Treatment of these same H118Y SUN1-expressing Huh7 cells with a CUL3-specific neddylation inhibitor, DI-1859 (Zhou *et al*, 2021), did not completely block CUL3 neddylation or restore IRS-1 expression, and accordingly its effect on insulin-stimulated AKT phosphorylation was modest (**Fig.S2**). While these data do not completely exclude CUL3-independent mechanisms by which SUN1 might regulate insulin signaling, we note that although other CRLs may be able to target IRS-1 for degradation *in vitro*, it appears that only CRL3 mediates IRS-1 degradation in hepatocytes *in vivo* (Chen *et al*., 2022). Taken together, these data support a model in which WT SUN1 maintains hepatic insulin signaling via control of CUL3 neddylation and preservation of cellular IRS-1 levels, while H118Y SUN1 is unable to bind to CUL3, leading to increased CUL3 neddylation, increased CRL3 activity, decreased IRS-1 levels, and blunted insulin signaling to AKT (**Fig.5B**).

**Fig. 5.**
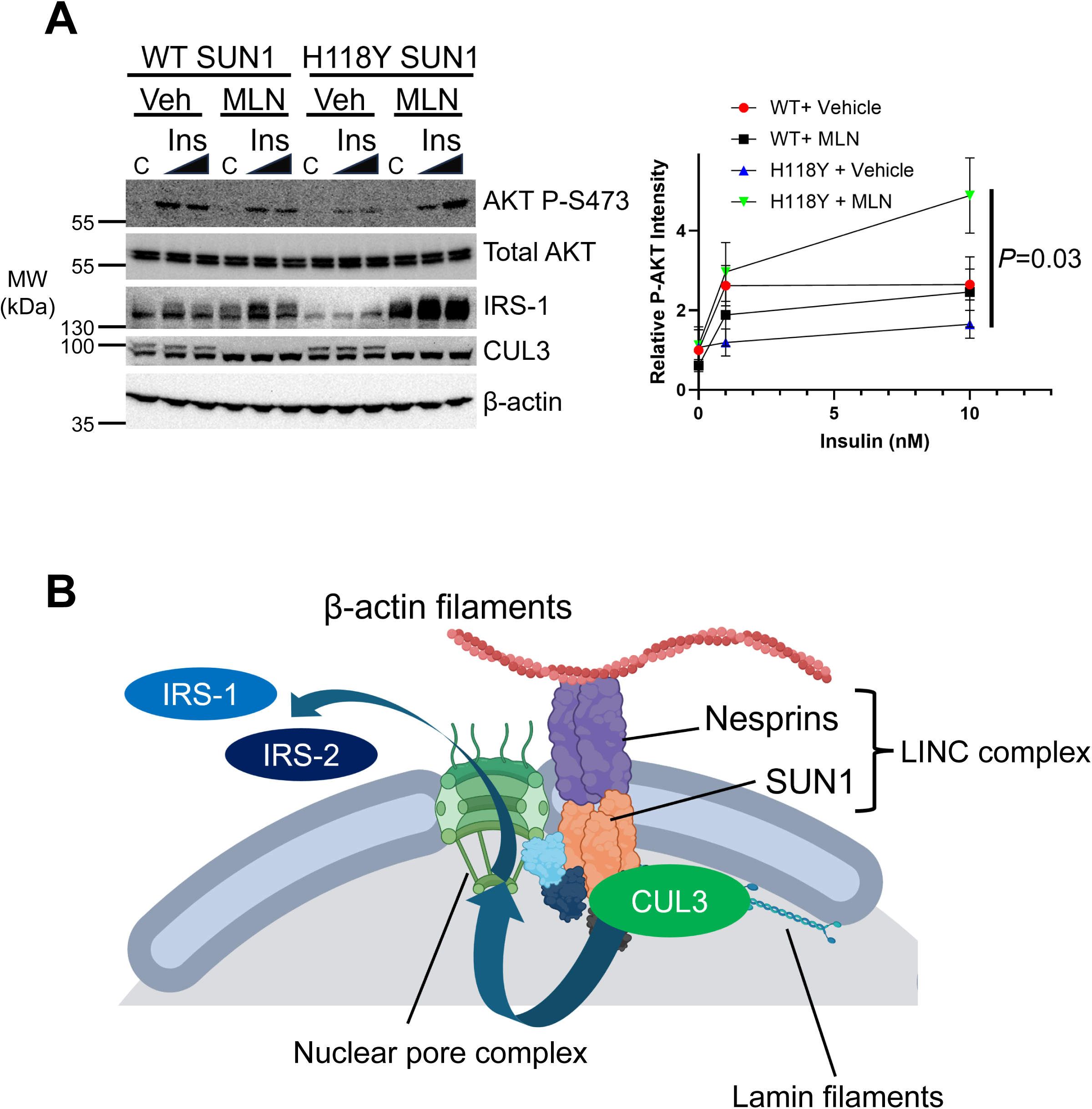
Inhibition of cullin neddylation restores insulin signaling in H118Y SUN1-expressing cells. *A*, Huh7 cells stably expressing SUN1-mCherry were treated with 1 or 10 nM human insulin for 20 minutes, with or without MLN4924 pre-treatment (1 µM, 16h), followed by lysis, SDS-PAGE, and immunoblot. C = control medium; Veh = vehicle (DMSO). Results shown are representative of, and quantitation shown at right was derived from, n=3 independent experiments. P-AKT intensity was relative to total AKT (ImageJ), with WT/vehicle control set to 1; *P*-value derived via ANOVA. *B*, proposed model of how wild-type SUN1 preserves cellular IRS protein levels by sequestration of inactive, non-neddylated CUL3; unbound CUL3 is available to become neddylated and target cytosolic IRS-1 and -2 for ubiquitination and degradation. BioRender was used under an institutional license to the University of Michigan in the production of part *B*.

## Discussion

We previously found that a common coding variant (H118Y) in *SUN1* positively associated with hepatic steatosis, histologic MASLD, and related metabolic traits, including several linked tightly to insulin resistance, in large population-based cohorts, and that ectopically expressed H118Y SUN1 blunted insulin signaling to AKT in human liver-derived cells; however, the molecular mechanisms by which SUN1 might regulate insulin signaling were unclear. Here we report that unbiased proteomics revealed a key SUN1-CUL3 physical interaction that maintains insulin signaling by restricting CUL3 neddylation and activation of CRL3-mediated ubiquitination and proteasomal degradation of downstream targets, including IRS-1. This interaction is impaired by the H118Y variant of SUN1, thus weakening this regulatory axis. Our data demonstrate that either loss of SUN1 function or the presence of H118Y SUN1 leads to aberrant CUL3 neddylation, IRS-1 degradation, and blunted insulin signaling to AKT in liver-derived Huh7 cells. Importantly, pharmacologic inhibition of CUL3 neddylation restored IRS-1 expression and insulin-stimulated AKT phosphorylation in H118Y SUN1-expressing cells, thus providing not only a potential mechanism for H118Y SUN1-driven insulin resistance but also a viable therapeutic approach for its reversal.

Two potentially surprising findings of our studies relate to the impacts of the neddylation inhibitors MLN4924 and DI-1859 on CUL3 neddylation and insulin signaling in Huh7 cells. First, it is somewhat surprising that MLN4924 has only a modest impact on IRS-1 levels and insulin signaling to AKT in WT SUN1-expressing cells, given that complete blockade of CUL3 neddylation is observed (Fig.5A). It is possible that compensatory mechanisms exist to limit IRS-1 expression and insulin signaling in WT SUN1-expressing cells, or that CRL3 activity is nearly maximally blocked even in the absence of MLN4924 in these cells. It will be important for future studies to address the underlying mechanism(s) of this modest effect of MLN4924 in the presence of WT SUN1 and whether it is observed *in vivo*, which could have significant implications for the impact of *SUN1* genotype on the efficacy of therapeutic neddylation inhibitors. Secondly, it is somewhat surprising that DI-1859 appears to have a more modest effect on both AKT phosphorylation and CUL3 neddylation than MLN4924, despite being previously reported to have a greater potency (Zhou *et al*., 2021). It is possible that this is a cell type-specific effect; it does not appear to be directly related to SUN1, as we observed persistent CUL3 neddylation in WT and H118Y SUN1-expressing and in parental Huh7 cells, at all tested concentrations of DI-1859, whereas MLN4924 completely blocked CUL3 neddylation at the same concentrations. This may or may not have significant implications for therapeutic cullin neddylation inhibition *in vivo*, since DI-1859 has been previously shown to have strong potency *in vivo*, but it may have significant implications for *in vitro* mechanistic studies that require full blockade of CUL3 neddylation.

The current study bridges two areas of emerging importance in MASLD and metabolic disease: the roles of inner nuclear membrane proteins and cullin-containing ubiquitin ligases. The role of inner nuclear membrane proteins in MASLD and metabolic disease has become clear in recent years, with a combination of animal studies and human genetics linking *TOR1AIP1*, *TOR1B*, and most recently *SUN1* to MASLD (Chen *et al*., 2023; Shin *et al*., 2019; Upadhyay *et al*., 2023). Among these, though, only *SUN1* appears to have a genetic and mechanistic link to insulin signaling and insulin resistance. The current study provides a potential explanation for why that might be the case: a unique SUN1 interaction with CUL3, a master regulator of protein stability and insulin signaling. The role of CUL3 in regulation of insulin signaling has recently become clear and is thought to occur via selective degradation of IRS-1 (and potentially IRS-2), which both act downstream of the insulin receptor to activate AKT phosphorylation (Chen *et al*., 2022; Gu *et al*., 2024; Gu *et al*, 2025). Either pharmacologic inhibition of cullin neddylation or hepatocyte-specific knockout of *CUL3* stabilizes IRS proteins and improves hepatic insulin sensitivity, although complete loss of CUL3 in hepatocytes led to systemic metabolic disturbances, including peripheral insulin resistance, which appeared to be due to uncontrolled activation of the NRF2 pathway and reductive stress (Gu *et al*., 2024). Our current study, rather than contradicting these findings, provides further clarity on the physiologic mechanisms by which CUL3 neddylation and activation are regulated. While complete loss of CUL3 in hepatocytes appears to be detrimental, restriction of inappropriate CUL3 neddylation is vital for the preservation of hepatic insulin signaling. This is important, because hepatic insulin resistance is thought to be a key feature of progressive, fibrotic MASLD and a determinant of both MASLD progression and cardiometabolic risk (Bo *et al*., 2024; Steinberg *et al*., 2025). Collectively, in the context of prior studies, our current findings underscore the dual role of CUL3 neddylation in regulation of insulin signaling and maintenance of systemic metabolic balance, making it a promising therapeutic target in insulin resistance and MASLD.

The current study has both strengths and limitations that should be acknowledged. Strengths of our study include a solid basis in our prior human genetic findings (Upadhyay *et al*., 2023), as well as robust and unbiased proteomic data, with confirmatory biochemical studies, derived from an experimental system with ectopic human SUN1 expressed at physiologic levels. Additionally, our data provide a novel and variant-specific mechanism by which CUL3 neddylation and insulin signaling are regulated in liver-derived cells. The major limitation of our study is that, while human genetic data support the role of SUN1 in protecting from MASLD and insulin resistance, and while the role of CUL3 in hepatic insulin signaling *in vivo* is already established, our data linking SUN1 to insulin signaling via CUL3 were derived *in vitro*; thus, although we expect future *in vivo* studies to corroborate our current findings, this will need to be confirmed empirically before these findings can be considered definitive.

In summary, our study has identified a novel mechanistic link between the inner nuclear membrane protein SUN1 and CUL3 neddylation that maintains insulin signaling to AKT and is impaired by the common MASLD and insulin resistance-linked *SUN1* variant H118Y. Pharmacologic blockade of cullin neddylation restored insulin signaling to AKT in H118Y SUN1-expressing cells, pointing toward a potential avenue for therapeutic targeting of insulin signaling in carriers of the common *SUN1* H118Y variant, although additional mechanistic studies – including *in vivo* studies – will be required to better understand the mechanisms and consequences of SUN1-mediated regulation of CUL3. Future *in vivo* studies should address whether the role of SUN1 in insulin signaling is entirely via CUL3 regulation, or whether CUL3-independent mechanisms may exist. Additionally, it will be important to determine whether cullin inhibition can improve insulin sensitivity in the presence of WT SUN1, or whether such an approach is likely to have limited utility in the absence of deregulated neddylation (due to H118Y SUN1 or another cause). A full mechanistic understanding of SUN1-mediated regulation of CUL3 will be critical in order to facilitate its therapeutic targeting in a way that is more precise than pan-neddylation or full blockade of CUL3 function, either of which may have detrimental metabolic consequences.

## Abbreviations

AP-MS: affinity purification-mass spectrometry
BCA: bicinchoninic acid
CRL: cullin RING ligase
CUL3: cullin-3
DMEM: Dulbecco’s Modified Eagle Medium
DTT: dithiothreitol
ECL: enhanced chemiluminescence
FBS: fetal bovine serum
FDR: false discovery rate
HEK: human embryonic kidney
IP: immunoprecipitation
IRS: insulin receptor substrate
LINC: Linker of Nucleoskeleton and Cytoskeleton
MASLD: metabolic dysfunction-associated steatotic liver disease
NEDD8: neural precursor cell expressed, developmentally down-regulated 8
PBS: phosphate-buffered saline
PCR: polymerase chain reaction
RING: really interesting new gene
RIPA: radioimmunoprecipitation assay
siRNA: small interfering ribonucleic acid
SUN1: Sad1 and unc84 domain containing 1
WT: wild-type.

## Acknowledgements

The authors gratefully acknowledge the Proteomics Resource Facility (RRID: SCR_026723) at the University of Michigan for their assistance with proteomic data acquisition and the University of Michigan Vector Core for assistance with lentivirus generation and production.

