## Supplementary Material for "The inner nuclear membrane protein SUN1 regulates cullin-3 neddylation to maintain insulin signaling"

**Table S1**. Top differential WT versus H118Y SUN1-interacting proteins (related to Figure 2).

| Protein | # of peptides | # of unique peptides | Ratio H118Y:WT | *P*-value |
|---|---|---|---|---|
| Cullin-3 | 4 | 4 | 0.01 | $1.55 \times 10^{-16}$ |
| SUMO4 | 2 | 1 | 8.1 | $1.55 \times 10^{-16}$ |
| Annexin A6 | 23 | 23 | 0.36 | $3.4 \times 10^{-15}$ |

### Figure S1

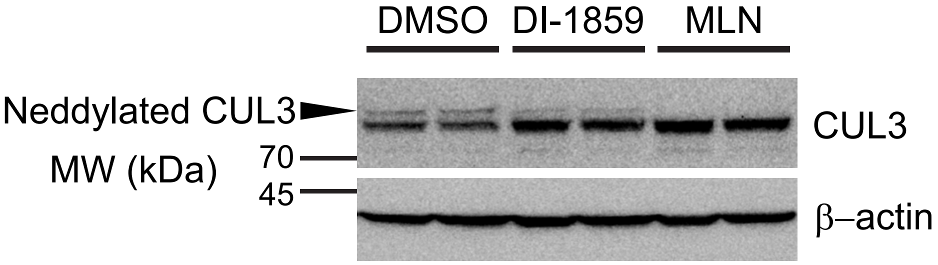


**Figure S1.** The slower-migrating CUL3 band represents neddylated CUL3. Huh7 cells were treated with DMSO (vehicle) or either 1 μM DI-1859 (CUL3-specific neddylation inhibitor, middle 2 lanes) or 1 μM MLN4924 (pan-neddylation inhibitor, last 2 lanes, shortened to "MLN") for 16h prior to lysis, SDS-PAGE, and immunoblot with anti-CUL3 antibody. Although a portion of CUL3 remained neddylated after DI-1859 treatment, no neddylated CUL3 could be detected after treatment with MLN4924. Results shown are representative of n=3 independent experiments.

### Figure S2

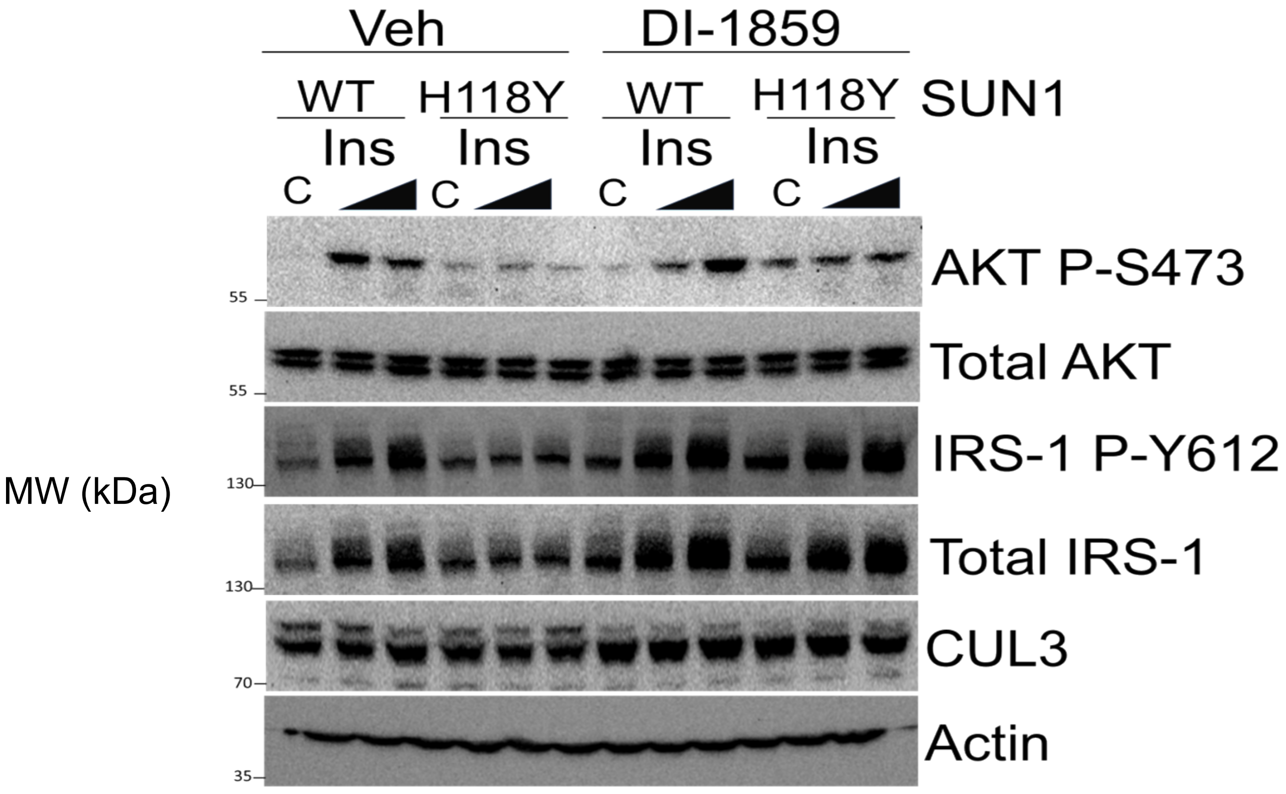


**Figure S2**. Partial rescue of insulin signaling to AKT in H118Y SUN1-expressing cells after treatment with DI-1859. Huh7 cells stably expressing SUN1-mCherry were treated with 1 or 10 nM human insulin for 20 minutes, with or without DI-1859 pre-treatment (1 µM, 16h), followed by lysis, SDS-PAGE, and immunoblot. C, control medium; Veh, vehicle (DMSO). Results shown are representative of n=3 independent experiments.
